# Designing a simple and efficient phage biocontainment system using the amber suppressor initiator tRNA

**DOI:** 10.1101/2024.07.29.605542

**Authors:** Pamela R. Tsoumbris, Russel M. Vincent, Paul R. Jaschke

## Abstract

Multidrug-resistant infections are becoming increasingly prevalent worldwide. One of the fastest emerging alternative and adjuvant therapies being proposed is phage therapy. Naturally-isolated phages are used in the vast majority of phage therapy treatments today. Engineered phages are being developed to enhance the effectiveness of phage therapy, but concerns over their potential escape remains a salient issue. To address this problem, we designed a biocontained phage system based on conditional replication using amber stop codon suppression. This system can be easily installed on any natural phage with a known genome sequence. To test the system, we mutated the start codons of three essential capsid genes in the phage ϕX174 to the amber stop codon (TAG). These phages were able to efficiently infect host cells expressing the amber initiator tRNA, which suppresses the amber stop codon and initiates translation at TAG stop codons. The amber phage mutants were also able to successfully infect host cells and reduce their population on solid agar and liquid culture but could not produce infectious particles in the absence of the amber initiator tRNA or complementing capsid gene. We did not detect any growth-inhibiting effects on *E. coli* strains known to lack a receptor for ϕX174, and we show that engineered phages have a limited propensity for reversion. The approach outlined here may be useful to control engineered phage replication in both the lab and clinic.

## Introduction

Broad-spectrum antibiotics are commonly used in medicine and agriculture to treat a wide range of bacterial infections. However, widespread administration is leading to the development and spread of multidrug-resistant (MDR) pathogenic bacteria. Consequently, patients infected with MDR pathogens have increased hospital stays [1, 2] and higher mortality rates [3]. The overuse of antibiotics also disrupts the microbiome [4] and may exacerbate chronic conditions from kidney disease to obesity [5].

One of the more promising avenues of treatment of MDR bacteria is the use of bacterial viruses called bacteriophages [6, 7]. Research into the therapeutic use of phages began over a century ago, but phage therapy, as it is now called, is becoming more prevalent as an alternative or adjuvant treatment for MDR bacterial infections [8, 9]. There are a growing number of reports outlining the successful treatment of MDR pathogens by phage therapy [10-16], as well as their ability to work synergistically with antibiotics [17-19].

Despite the impressive and growing track record of phage therapy, the treatment has limitations [9, 20] that prevent it from being administered in a wider context. The promise of low-cost, broad-spectrum phage therapy is constrained by the natural features of phages, such as narrow host ranges, rapid development of phage resistance, concern over off-target effects on the human microbiome, and phage-mediated horizontal gene transfer [21, 22]. These obstacles to broader adoption may be overcome by employing the tools of synthetic biology to re-engineer phages [23-27], broadening or adapting host range [28, 29], making them more resistant to the antiphage defence systems of microbes [30], improving their killing efficacy through the addition of antimicrobial payloads [31-33], and increasing the range of phages available by the removal of lysogeny genes [34].

One potential barrier to the effective implementation of engineered phages is associated regulatory concerns and hurdles regarding their uncontrolled release into patients and the environment [35]. One method of addressing these concerns is through engineered genetic biocontainment mechanisms that aim to limit the spread of the phage beyond its intended location. In this context, genetic biocontainment involves manipulating microbe genomes to prevent the engineered organism’s unintended proliferation, spread, or persistence outside the designated environment [36, 37]. Biocontainment strategies have been devised for different microorganisms using genetic code modification [38, 39], auxotrophy [40, 41], kill switches [42, 43], dependence on unnatural amino acids [44, 45] and combinations of different systems [46, 47]. While effective, these methods are not easily scalable, and many have organism-specific requirements. Furthermore, many of these biocontainment methods are not suitable for use in phages due to inherent aspects of their design.

In this work, we present a simple and effective phage biocontainment system that requires minimal knowledge of the phage and does not change the phage genome length. The system is based on engineering phage replication to be dependent on the amber suppressor initiator tRNA [48] that initiates orthogonal translation at the UAG stop codon [49, 50]. We show that the system is easily implemented and prevents phage replication in the absence of the amber initiator tRNA. Furthermore, we show that non-replicating phage can infect and lyse target bacterial strains while not affecting the growth of non-target strains.

## Methods

### Bacterial strains, plasmids and growth conditions

The *E. coli* strains employed in this study included M1402 [51], MG1655, Nissle 1917, C900 (a gift from Bentley Fane; originally created by Ry Young [52]), and NCTC122 (hereby referred to as C122; National Collection of Type Cultures, Public Health England). All propagation and expression of amber initiated ϕX174 (aiϕX174) mutants used C122(pULTRA-*metYp1p2-metY*(CUA)), which placed the amber initiator i-tRNA_CUA_ under the native constitutive promoter *metYp1p2 [53]. E. coli C122* strains expressing either the F, G, or H gene from a rhamnose-inducible promoter were also created by transforming them with one of the medium copy number pJ804-gene F [23], pJ804-gene G, or pJ804-gene H plasmids. Expression of the F, G or H genes from these plasmids was induced with 1 mM rhamnose. All bacterial strains were grown in Lysogeny Broth (Miller) (10 g/L Tryptone, 10 g/L NaCl, 5 g/L Yeast Extract, dissolved in deionized water 18 MΩ-cm) supplemented with 2 mM CaCl_2_ hereby referred to as phage LB (PLB) [54]. PLB was supplemented with spectinomycin (50 μg/mL) or carbenicillin (50 μg/mL) where appropriate. Unless otherwise stated, all overnight bacterial cultures were grown for 18 hours until stationary phase was reached in fresh 15 mL centrifuge tubes at 37°C, rotating orbitally at 250 RCF at 25 mm diameter. All overnight cultures used were <48 hours old.

### Creating phage mutants

All aiϕX174 mutants were designed to match the wild-type ϕX174 sequence from the original Sanger sequence [55] (Genbank No. NC_001422.1), which differs from the commercially available preparations by nine bases [56] [23, 57]. The ϕX174 genome was divided into four sections labelled F1, F2, F3a/F3b and F4 (Fig. S1 and Table S1). Fragments F1, F3a and F3b were synthesized (IDT) with a 2-base change at the ATG start codons of the F, G and H genes to a TAG amber stop codon, and were subsequently used to create F, G and H amber mutants (*Fam, Gam* and *Ham*, respectively). A WT ϕX174 plaque was suspended in sterile water and used to generate fragments F2 and F4 via high-fidelity PCR (Table S2). PCR products were then subjected to DpnI digestion (1 hr/37°C), followed by 20 min/ 80°C incubation, then cleaned up (Monarch PCR Clean Up Kit, NEB). Fragments containing either the ATG start codon or TAG start codon were combined in a 1:1 ratio for a Gibson Assembly [58] (Table S3) and incubated at 50°C for 1 hour. The assembled constructs were transformed into C122-*metYp1p2-metY*(CUA) cells made competent via Mix & Go (Zymo Research, cat. #T3007). PLB was added to a total of 1 mL, and the cells were heat-shocked at 42°C for 30 seconds. The infected cells were then incubated at 37°C and agitated at 250 rpm for 1 hour. Plaque assays were performed on C122-*metYp1p2-metY*(CUA) and resulting plaques were picked for PCR screening and sequence confirmation.

### Phage strains and growth conditions

For all ϕX174 variants used in this study the Phage-On-Tap protocol [59] was adapted for propagation and purification. From a fresh overnight culture, C122(pULTRA-*metYp1p2-metY*(CUA)) was inoculated 1:100 into 10 mL PLB grown in a 50 mL centrifuge tube and incubated at 37°C with continuous orbital shaking at 250 rpm (25 mm diameter). Once the starter culture reached exponential phase (OD_600_ = 0.4 – 0.5), it was infected with a plaque isolated from a plate of phages suspended in 1 mL SM (50 mM Tris-HCl (pH 7.5), 8 mM MgSO_4_, 100 mM NaCl, dissolved in deionized water 18 MΩ-cm) buffer and incubated for approximately 2 hours or until the culture was cleared. The resulting phage lysate was passed through a 0.2 μm filter to remove remaining bacteria and debris and stored overnight at 4°C. This process was then repeated using a total volume of 100 mL in a 250 mL baffled glass flask. Instead of infecting exponential bacteria with individual plaques, the phage lysate from the starter culture was used to infect the bacterial culture. Once OD_600_ < 0.2 (∼ 3 hours), 0.1 volumes of chloroform and 1 M NaCl were added, and the resulting resuspension was centrifuged at 4 000 × g for 10 minutes to remove bacterial debris, chloroform, and uninfected cells. The supernatant was passed through 0.2 μm filters to remove any remaining debris, then subjected to concentration using Amicon Ultra-15 Centrifugal Filter units (Merck cat. # Z740208-8EA) with a 100 kDa molecular weight cut-off. Once the volume of phage was reduced to ∼5 mL, 10 mL of SM buffer was added to the filters and centrifuged at 4 000 × g to wash the phage until ∼1 mL volume remained. The concentrated phage lysate was then titred via plaque assays, as described below.

The biocontainment capacity of each aiϕX174 was tested on several strains with varying degrees of susceptibility. Susceptibility of C122, M1402, MG1655, and Nissle1917 to WT ϕX174 were first determined via the plaque assay method. MG1655 and Nissle1917 were chosen as they are not susceptible to ϕX174 infection and were therefore determined to be adequate controls due to their laboratory context and potential as a probiotic, respectively. To test the capacity of phages to lyse at varying multiplicities of infection (MOIs), ten-fold dilutions were made of WT ϕX174, *Fam, Gam* and *Ham* phages, starting with concentrations as high as 1×10 PFU/mL. A spot assay was then conducted, as described below, using the phage dilutions. This was also completed for C122 strains harbouring plasmid expressing the F, G and H genes to identify potential polar effects from the genome modifications.

The minimum inhibitory concentration (MIC) of the aiϕX174 infection was also tested on the above strains via a liquid culture, using *Fam* as a representative of the variants. To achieve this, an overnight culture was diluted 1:100 in fresh PLB and grown at 37°C, rotating orbitally at 250 rpm until mid-log phase was reached. Cultures were then diluted to ∼100 cells in a 96-well clear-well, round side, flat bottom microplate. *Fam* was added in varying concentrations to achieve MOIs 1, 10, 1,000 and 10,000, achieving a final volume of 200 µL. Cultures were then covered with Breathe Easier sealing membranes (cat. #Z763624) and incubated at 37°C with double orbital shaking at 287 rpm (3 mm diameter). The OD_600_ was monitored every 10 minutes using a plate reader (Biotek Synergy H1).

### Plaque and Spot Assays

Plaque assays were performed using 1.5% LB agar plates and 0.7% PLB top agar. To perform the plaque assays, 100 μL of fresh overnight host culture was suspended in 3 mL of molten top agar (50°C) and then infected with 10 μL of phage. Plates were left to dry at room temperature for 5 minutes. Infected plates were incubated overnight at 37°C and corresponding plates were imaged using the Gel Doc Image Lab software (BioRad). Plaque forming units per mL (PFU/mL) were calculated, and plaque diameters were measured from images using the Plaque Size Tool [60]. The generated data was then analyzed using the Mann-Whitney U test for significance using R (v4.3.1). To perform a spot assay, plates were set up as previously described for plaque assays without the addition of phage. Once the top agar had solidified, plates were incubated at 37°C for 1 hour. Phage stocks were serially diluted ten-fold in SM buffer and 3 µL then added to the top layer. Plates were left to dry for 10 minutes or until there was no longer liquid visible on top of the agar, and then incubated and imaged as above.

### Cell Lysis Curves

Lysis curves were set up in triplicate cultures for the aiϕX174 variants. Bacterial host strains were grown in PLB to OD_600_ 0.5, then diluted 1:100 into 96-well clear-well, round side, flat bottom microplate to a final volume of 200 μL. Phages were added to each sample at MOI = 0.01. Microplates were covered with Breathe Easier sealing membranes (cat. #Z763624) and incubated at 37°C with double orbital shaking at 283 rpm (3 mm diameter), with OD_600_ measured in 5-minute intervals (Biotek Synergy H1 plate reader). Similarly, cell lysis curves were performed for the *Fam* variant to test lytic efficacy of the mutant phage at high MOIs. Triplicate bacterial host strains were grown in PLB to OD_600_ 0.5, then diluted to a final volume of 200 μL in a 96-well clear-well, round side, flat bottom microplate. Each well contained approximately 100 cells. Phages were added to each sample at an estimated MOI of 1, 10, 100, 1000, 10 000. Microplates were covered with Breathe Easier sealing membranes (cat. #Z763624) and incubated at 37°C with double orbital shaking at 283 rpm (3 mm diameter), with OD_600_ measured for 16 hours.

### Measuring Host Resistance and Fam revertant frequency

Host resistance and phage revertant frequency were determined by performing plaque assays on the C122 host with *Fam* phage lysate. Wild-type revertant phage would be expected to form plaques on the C122 strain, while *Fam* mutants would only produce plaques in the presence of the i-tRNA. Additionally, we tested for the presence of phage-resistant hosts in treated samples by spreading 100 µL from the MOI=10,000 infected cultures after 18 hours of incubation at 37 °C on LB (Miller) agar plates. Plates were incubated at 37°C overnight (18 hours), and resulting colonies were counted. These colonies were then inoculated into PLB, grown overnight (18 hours) at 37°C with agitation of 200 rpm, and subsequently used to determine phage-sensitivity to WT ϕX174.

### Results

To create and test a phage biocontainment system based on amber stop codon suppression, we first created an *E. coli* strain capable of expressing the amber suppressor initiator tRNA (i-tRNA2 ^fMet^ _CUA_). Previously, we have shown that i-tRNA_2^fMet^ CUA_ can efficiently initiate translation from UAG start codons and that it does not have any detectable off-target translation initiation within *E. coli* [49, 50]. We created a new version of the plasmid we previously used by swapping the tacI promoter for the endogenous *metYp1p*2 promoter to make it independent of exogenously added inducer (Fig. 1A). This new plasmid pULTRA-*metYp1p2*(*metYCUA*) constitutively and stably expresses i-tRNA_2 ^fMet^ CUA_ [61] under regular lab growth conditions [62].

**Fig. 1.**
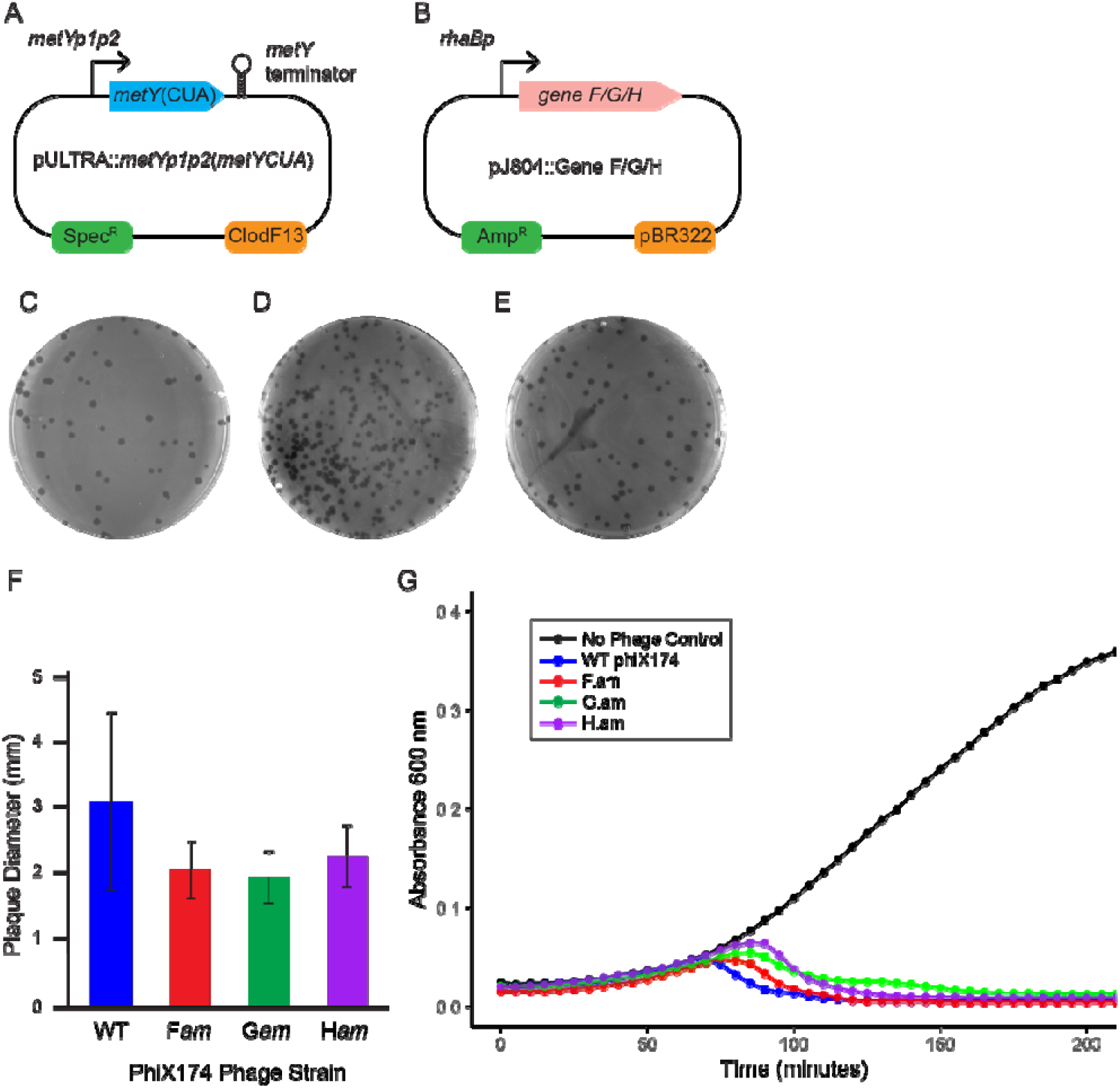
Design and construction of aiϕX174 phages and resulting plaque phenotype. (A) Map of the medium-copy pULTRA-*metYp1p2*-*metY*(CUA) plasmid. The native constitutive *metYp1p2* promoter controls the expression of wild-type initiator tRNA. (B) Map of medium-copy pJ804 plasmid containing ϕX174 genes F, G, or H driven by rhamnose-activated *rhaBp* promoter. (C) Fam, (D) *Gam*, (E) *Ham* phage plaques on MP strain. (F) Plaque sizes of aiϕX174 phages. (G) Lysis curve of WT ϕX174 and aiϕX174 phage on the MP strain. Cells were infected at an MOI of 0.01 (n=3), with line shading representing one standard deviation.

Next, we recoded the canonical ATG start codon of ϕX174 capsid genes F, G and H to the amber TAG stop codon, creating *Fam, Gam*, and *Ham* mutant ϕX174 phage, which collectively we term aiϕX174 for amber initiator ϕX174. We then transfected the mutant DNA into the C122(pULTRA-*metYp1p2*(*metYCUA*)) host strain (called hereafter MP). Resulting plaques (Figs. 1C, D, and E) were measured using Plaque Size Tool [60]. The average plaque diameters of *Fam* (2.1 ± 0.4 mm), *Gam* (2.0 ± 0.4 mm) and *Ham* (2.3 ± 0.5 mm) were all slightly smaller than WT ϕX174 (3.1 ± 1.3 mm) (Fig. 1F). Each of the aiϕX174 also showed less variation in plaque size (20-21%) compared to WT (43%). Lysis curves of the aiϕX174 phages on the MP strain showed that they had a slightly slower lysis time than WT ϕX174, although by 2 hours post-infection all phages showed a similar degree of lysis (Fig. 1G).

### Amber mutant phages cause lysis without plaque formation in strains lacking the amber initiator Trna

To assess the ability of the aiϕX174 phages to lyse host cells, we measured phage infection characteristics in five different strains of *E. coli*. The strain C122 was used as the regular host, M1402 as a host that displays lower efficiency of plating compared to C122, Nissle1917 and MG1655 as negative control strains that are non-susceptible to ϕX174, and C900 *slyD* mutant strain which is resistant to ϕX174 lysis via disruption of the lysis protein E maturation pathway. Additionally, we included the MP strain as a positive control for unhindered aiϕX174 mutant phage replication. We also constructed three plasmids (pJ804-Gene F, pJ804-Gene G, and pJ804-Gene H) that could express ϕX174 proteins F, G, or H as complementation controls for the individual *Fam, Gam*, and *Ham* phages (Fig. 1B).

We tested the panel of *E. coli* strains with varying concentrations of WT ϕX174, *Fam, Gam*, and *Ham* phages by the spot assay method on agar plates. The results of this experiment showed complete cell lysis for WT ϕX174 against i-tRNA-expressing strain MP and C122 (Figs. 2A and B). Lysis was also seen for C122(pJ804-Gene F), C122(pJ804-Gene G), and C122(pJ804-Gene H) strains expressing the ϕX174 F, G, and H proteins, respectively (Figs. 2G, H, and I). As expected, no growth inhibition of any kind was seen for WT ϕX174 infections of Nissle1917 or MG1655, which lack ϕX174 receptors (Figs. 2E and F). A small amount of growth inhibition was seen for the C900 strain at the highest phage concentration spots, which could be explained by a slight metabolic burden that ϕX174 infection placed on the cells (Fig. 2D).

**Fig. 2.**
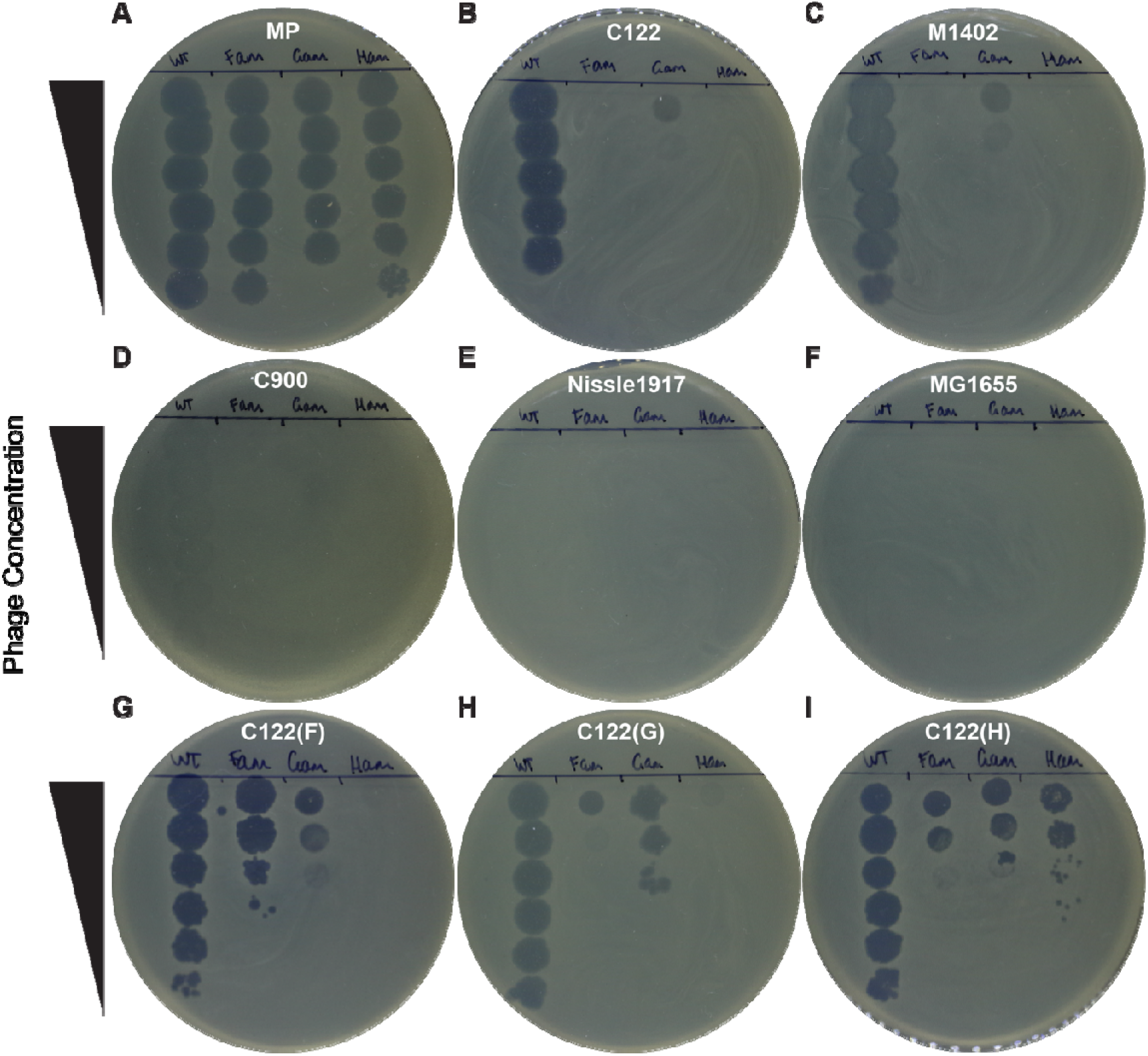
Biocontainment of aiϕX174 on solid media. Spot assays of WT ϕX174, *Fam, Gam* and *Ham* phages on different *E. coli* strains. (A) MP, (B) C122, (C) M1402, (D) lysis resistant C900 strain, (E) non-susceptible Nissle1917, (F) non-susceptible MG1655, (G-I) C122 expressing the ϕX174 F, G or H gene in the absence of amber initiator tRNA expressing plasmid. Highest concentration spot is 10^6^ plaque forming units, as measured on C122 for WT ϕX174 or MP strain for *Fam, Gam*, and *Ham* phages. To induce pJ804-Gene F/G/H plasmids, 1 mM of rhamnose was added to the top agar.

The *Fam, Gam* and *Ham* mutant phages all showed vigorous lysis when infecting the MP control strain containing the amber initiator tRNA (Fig. 2A). The *Fam* and *Ham* phages did not show any discernible spot of growth inhibition or lysis on C122 or M1402 strains in the absence of amber initiator tRNA expression (Figs. 2B and C). These phages also did not show any lysis against the non-permissive strains Nissle1917 or MG1655 (Figs. 2E and F) or growth inhibition against the C900 strain (Fig. 2D). By contrast, the *Fam* phage did show zones of bacterial growth inhibition and lysis, but no plaques, on C122 strains expressing the ϕX174 G or H proteins (Figs. 2G and I). Similarly, the *Ham* phage showed a slight amount of growth inhibition on the C122(pJ804-Gene G) strain (Fig. 2H).

In contrast to the *Fam* and *Ham* phage, the *Gam* phage showed obvious zones of bacterial growth inhibition and lysis at the two highest phage concentrations on the native C122 host, as well as the M1402 strain (Figs. 2B and C). Like the other amber mutant phage, *Gam* did not show any effect on the Nissle1917 or MG1655 strains (Figs. 2E and F) nor any effect on the C900 strain (Fig. 2D). Moreover, *Gam* showed enhanced growth inhibition and lysis, but not plaque formation, when infecting C122(pJ804-Gene F) and C122(pJ804-Gene H) strains (Fig. 2G and I).

Infection of the *Fam, Gam*, and *Ham* mutant phages against strains capable of complementing their mutated gene showed a different pattern. Each phage showed zones of lysis in the most concentrated phage spot, while at the least concentrated spot, distinct plaques could be seen (Figs. 2G, H, and I). This result showed that lysis in these strains was due to the production of infectious virions and phage replication, as opposed to phage lysis without replication. The efficiency of plating of the *Fam, Gam*, and *Ham* phages was lower in the complementing C122(pJ804-Gene F/G/H) strains (Figs. 2G, H, and I) than on the MP strain (Fig. 2A).

### Assessment of Fam phage for growth inhibition and revertant frequency in liquid culture

To further characterize the ability of aiϕX174 phages to inhibit bacterial growth in liquid culture, we used the *Fam* phage as a representative. We infected cultures of C122, M1402, K-12, and Nissle1917 with *Fam* phage at six different MOIs and measured an OD_600_ endpoint after 18 h incubation. The results of this experiment showed that *Fam* was able to completely inhibit C122 growth at MOI=10,000 and relative phage:host concentrations of 5 × 10^6^ PFU/mL to 5 × 10^2^ CFU/mL, whereas M1402 growth was only partially inhibited under these conditions (Fig. 3). This is consistent with the lower efficiency of plating of ϕX174 on M1402. C122 growth was partially inhibited at MOI=1,000, but M1402 growth was not inhibited at this MOI (Fig. 3). As expected, the growth of control strains K-12 and Nissle1917 was not inhibited at any MOI.

**Fig. 3.**
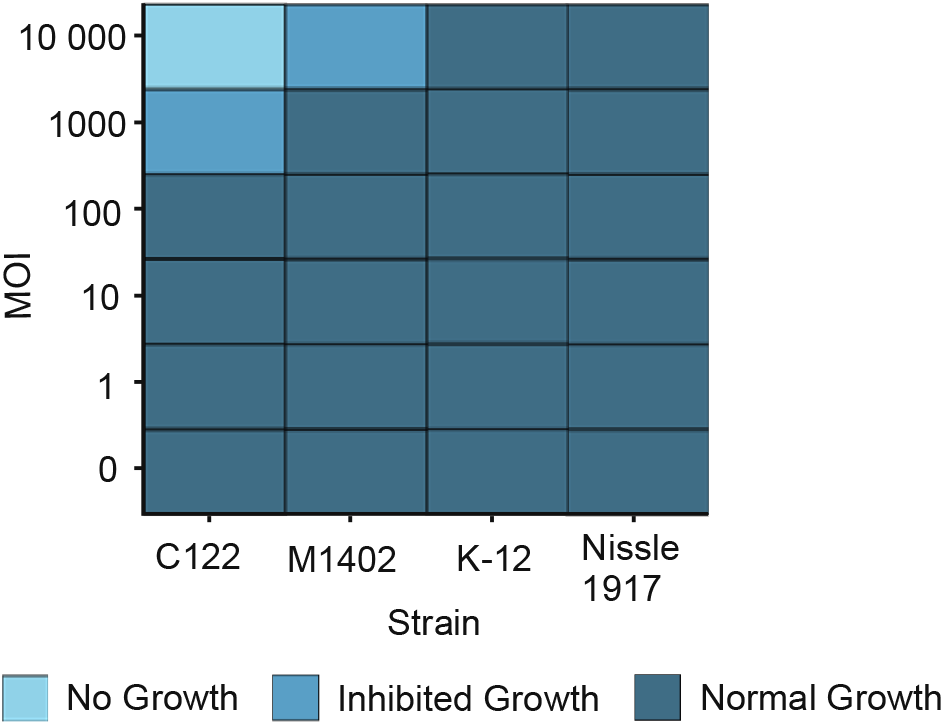
*Fam* phage shows growth inhibition at high multiplicity of infection in the native host and a low efficiency of plating strain. Infection of native C122, low efficiency of plating strain M1402, and non-permissive MG1655 and Nissle1917 strains. An average of 100 cells at a concentration of 5 × 10^2^ CFU/mL were infected with *Fam* across MOIs = 0 – 10^6^ 000 at phage concentrations ranging from 0 – 10 PFU/mL. Infected cultures were incubated with 287 rpm shaking at 37°C for 18 hours.

To determine whether there were any free *Fam* phage remaining after the 18 h incubation, we mixed samples of the MOI = 10,000 condition with the MP strain and plated using the double agar overlay method. We detected no plaques from this experiment, showing that all the input *Fam* phages were consumed during the 18 h incubation. To detect if any WT revertant phage were produced during the 18 h incubation, we mixed samples of the MOI = 10,000 condition with C122 host strain but were again unable to recover any plaques. We plated MOI = 10,000 condition endpoint cultures onto agar plates to determine the frequency of the development of phage resistance in the C122 host strain. We recovered six colony forming units from the liquid lysis curves over four biological replicates. These colonies appeared smaller than the initial parental C122 strain with a slight glossy sheen. After overnight growth in liquid media the cultures showed notable differences in appearance, with all displaying cell clumping and settling to the bottom of the tube rather than dispersion throughout the media as is seen in the ancestral C122 strain. Agitation caused cultures to disperse, but aggregation and settling quickly occurred afterwards. Infecting these cultures with WT ϕX174 at MOI = 1 showed no lysis over 18 hours, strongly indicating phage resistance. We estimated C122 phage resistance frequency in this experiment to be ≤ 2 × 10^−7^.

## Discussion

In this work we have outlined the development and characterization of a simple method to create biocontained bacteriophage by engineering their reliance on the amber initiator tRNA for essential gene expression and replication. This method only requires the identification of the start codon of at least one essential gene, and the ability to deploy a plasmid expressing the amber initiator tRNA into the phage host strain, or integration of the amber initiator tRNA into the host genome [63]. Currently, the system is optimized for *E. coli* but could be adapted for other species by modifying the essential characteristics of the device (promoter, initiator tRNA backbone, plasmid origin of replication) to accommodate the other species. The initiator tRNA is perfectly conserved across all three Domains of life, and amber initiator mutants have been shown to function in Gram-positive bacterium *Mycobacterium smegmatis* [64], as well as in human cells [65].

An advantage of this method over other alternative biocontainment methods is that changing the start codon of a gene from ATG to TAG is unlikely to disrupt any intragenic promoters in the affected gene sequence [66, 67]. Furthermore, although the removal of entire essential genes from phages has been demonstrated to work efficiently [68], the method requires prior knowledge of the gene function and overall phage biology to ensure viability is not compromised. This creates various obstacles to the use of novel phages, which may contain a large proportion of genes with poorly defined functions [69-72]. Furthermore, the removal of genes may alter viability as some phages are extremely sensitive to genome shortening [73]. Additionally, there is the potential for disruption to essential intragenic regulatory sequences, impacting adjacent gene expression [67, 74].

Limitations of the work presented here include its inability to effectively target overlapping genes [75]. A large number of overlapping genes in prokaryotes and viruses are 1-bp or 4-bp start-stop codon overlaps [76-78], that would be disrupted by changing the start codon from ATG to TAG.

There is also the potential for mutational reversion from the amber stop codon TAG to a canonical start codon ATG in the gene of interest. This change requires two simultaneous transversion mutations at an adjacent site on one genome. It is also possible that a single transversion mutation (e.g. GTG or TTG) or a single transition mutation (e.g. CTG) could result in a weak start codon [79, 80] at first, and then a second mutational event results in the canonical ATG start codon. Given that previous estimates of the ϕX174 mutation rate are relatively high [81], the fact that we did not observe this happening in any of our tests was surprising. One possibility is that the mutational frequency is not uniform across the entire ϕX174 genome. This theory is supported by the fact that the most common mutations are found in either the F or H genes, regardless of the host ϕX174 has infected [82]. Propagation of the aiϕX174 phages in a strain overproducing the amber initiator tRNA seems to result in very little selective pressure for a mutation in the start codon of the gene controlled by the engineered tRNA and our reversion estimate frequency (≤ 10^−7^) seems to support this notion.

While the aiϕX174 mutants appeared to replicate in the presence of the amber initiator tRNA in a very comparable manner to the WT ϕX174, we did see small plaque size reductions and lengthened lysis times in the MP strain (Figs. 1C - E). It is possible that increasing the strength of the amber initiator promoter, placing it on a higher-copy plasmid, or supplying additional auxiliary genes necessary for production of the initiator tRNA could increase the replication efficiency of the biocontained phage from the producer strain [83, 84]. The MP strain did outperform simple complementation with the full-length gene expressed from a plasmid (C122(pJ804-Gene F/G/H) strains), indicating that the pULTRA-*metYp1p2*-*metY*(*CUA*) plasmid is already highly efficient (Fig. 2).

In this work we observed that some aiϕX174 mutant phage performed better on agar plate spot tests than others (Fig. 2), pointing to some target-specific effects that may need to be taken into account when creating phage using this system. Similarly, a large excess of non-replicating phage seem to be necessary to lyse all target bacteria (Fig. 3), especially if the strain is infected poorly by the phage, as was seen for M1402. Reassuringly, the *Fam* phage was shown to be highly specific for strains containing receptors and did not affect the growth of non-target strains even at very high MOIs (Fig. 3).

The non-replicating phage produced by this method cannot expand its population exponentially to catch growing bacterial hosts as a replicating phage can. Therefore, the doses of non-replicating phage delivered in a therapeutic setting would need to be significantly higher or more frequently dosed to account for this difference. Phage therapy currently requires administration of a 1 × 10^9^ PFU/dose at least once a day, often twice per day for a minimum of one week [10, 13, 15, 16, 85, 86]. As such, a full weeklong course of phage therapy would result in the total administration of 7 × 10^9^ – 3 × 10^10^ PFU. If we assumed that a non-replicating phage treatment, as based on our results here, would require at least 1,000-fold more phage per dose due to their lack of expansion in vivo, then that would require approximately 10^12^ – 10^13^ PFU per therapy course. Although these values may seem large, they are actually fairly modest if compared to an antibiotic biologic such as bezlotoxumab, which requires ∼3 × 10^18^ molecules per treatment course [87] and costs ∼$2,000 USD per course.

On a cost basis, phage production at scale is relatively inexpensive per dose, but is significantly more expensive if produced in small batches as is currently required. For example, a single batch of 1,000 GMP-grade phage doses (∼30 treatment courses) costs ∼$40,000 USD to produce [88], or ∼$1,333 per patient [88]. In reality though, often only a few patients can be treated with these phage, driving the cost per patient significantly up. In this paper, we have provided a simple methodology to create biocontained phage so that engineered phage may be more widely used in phage therapy. The development of engineered phages with enhanced features that make them more broadly useful against a range of bacterial hosts would help drive this cost back down, as entire phage batches could be used.

## Acknowledgements

We recognize that this research was conducted on the traditional lands of the Wallumattagal clan of the Dharug nation. The authors thank all members of the laboratory for their contributions and support.

## Funding

This work was supported by NHMRC Ideas Grant 2019/GNT1185399. PRT and RMV were supported by the Macquarie University Research Excellence Ph.D. scholarship (MQRES).

## Author Contributions

All authors contributed to the conceptualization, design and funding acquisition of this study. Formal analysis and visualization was performed by Pamela R. Tsoumbris and Paul R. Jaschke. Investigation and methodology was conducted by Pamela R. Tsoumbris and Russel M. Vincent. Validation was performed by Pamela R. Tsoumbris and Paul R. Jaschke. Project administration and supervision was conducted by Paul R. Jaschke. The original draft of the manuscript was written by Pamela R. Tsoumbris and Paul R. Jaschke. All authors read and approved the final version of the manuscript.

## Conflicts of Interest

The authors declare no conflict of interest.

## Data availability

The data used and/or analyzed in this study are available from the corresponding author upon reasonable request.

## Notes

### Competing Interest Statement

The authors have declared no competing interest.

