## Supplementary Information for "Designing a simple and efficient phage biocontainment system using the amber suppressor initiator tRNA"

**CONTENTS**

SUPPORTING FIGURES

Figure S1: aiφX174 genome as designated by individual fragments.

SUPPORTING TABLES

Table S1: aiφX174 fragment sequences.

Table S2: Primers used to amplify WT φX174 backbone for fragments F1 and F4.

Table S3: Primers used for Gibson Assembly of fragments.

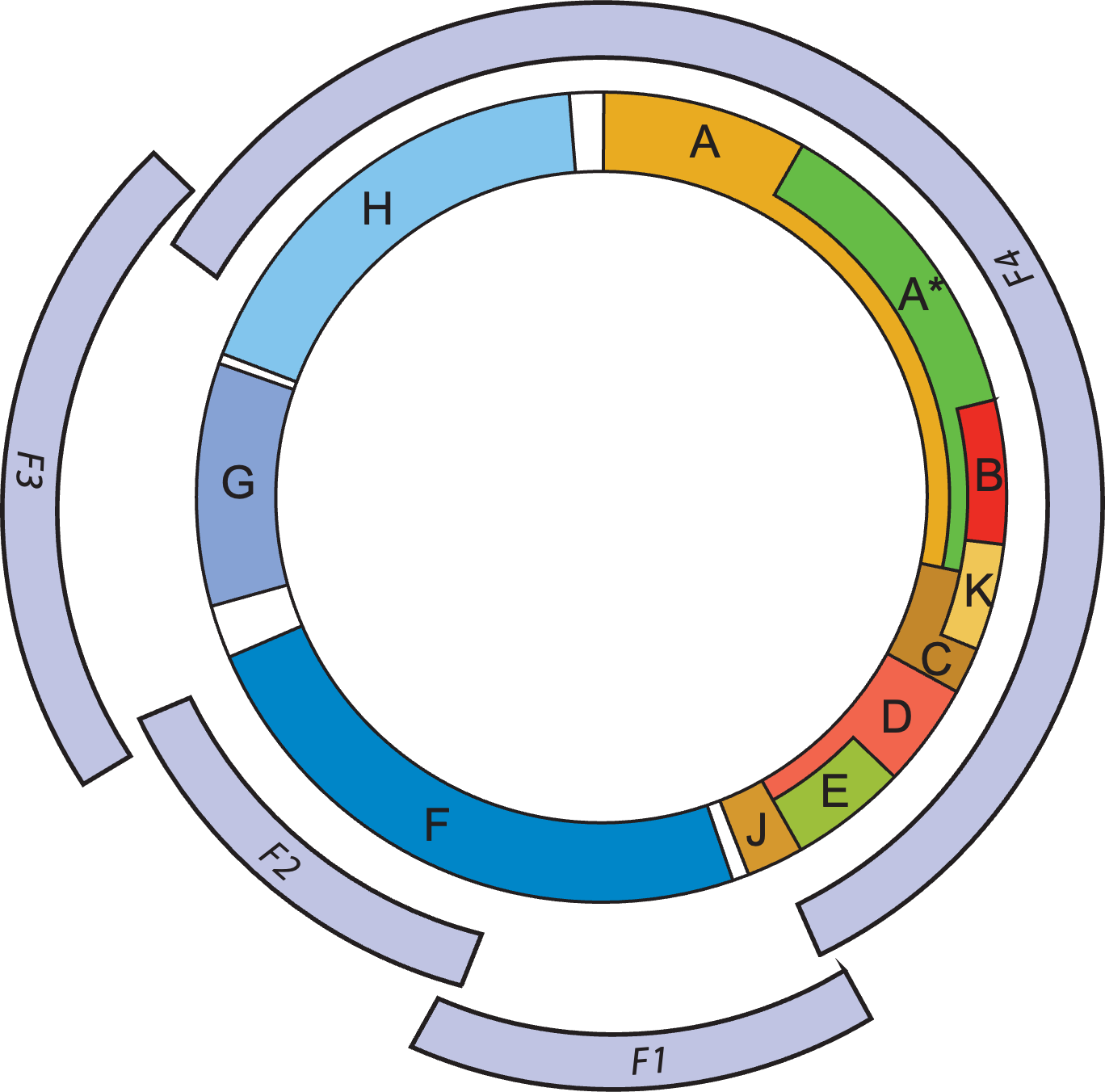

**Figure S1. The phiX174 genome divided into fragments to allow for Gibson Assembly.** The F start codon can be found in F1. The G and H start codons can be found in F3.

**Table S1. Sequences of aiφX174 fragments.** Highlighted in bold are the start sites of the F, G and H genes corresponding to each of their respective fragments.

| **Fragment** | **Sequence** |
| --- | --- |
| F1 (WT) | gtctttggtatgtaggtggtcaacaattttaattgcaggggcttcggccccttacttgaggataaatt**atg**tctaatattcaaactggcgccgagcgtatgccgcatgacctttcccatcttggcttccttgctggtcagattggtcgtcttattaccatttcaactactccggttatcgctggcgactccttcgagatggacgccgttggcgctctccgtctttctccattgcgtcgtggccttgctattgactctactgtagacatttttactttttatgtccctcatcgtcacgtttatggtgaacagtggattaagttcatgaaggatggtgttaatgccactcctctcccgactgttaacactactggttatattgaccatgccgcttttcttggcacgattaaccctgataccaataaaatccctaagcatttgtttcagggttatttgaatatctataacaactattttaaagcgccgtggatgcctgaccgtaccgaggctaaccctaatgagcttaatcaagatgatgctcgttatggtttccgttgctgccat |
| F1 (TAG) | gtctttggtatgtaggtggtcaacaattttaattgcaggggcttcggccccttacttgaggataaatt**tag**tctaatattcaaactggcgccgagcgtatgccgcatgacctttcccatcttggcttccttgctggtcagattggtcgtcttattaccatttcaactactccggttatcgctggcgactccttcgagatggacgccgttggcgctctccgtctttctccattgcgtcgtggccttgctattgactctactgtagacatttttactttttatgtccctcatcgtcacgtttatggtgaacagtggattaagttcatgaaggatggtgttaatgccactcctctcccgactgttaacactactggttatattgaccatgccgcttttcttggcacgattaaccctgataccaataaaatccctaagcatttgtttcagggttatttgaatatctataacaactattttaaagcgccgtggatgcctgaccgtaccgaggctaaccctaatgagcttaatcaagatgatgctcgttatggtttccgttgctgccat |
| F2 | gaggctaaccctaatgagcttaatcaagatgatgctcgttatggtttccgttgctgccatctcaaaaacatttggactgctccgcttcctcctgagactgagctttctcgccaaatgacgacttctaccacatctattgacattatgggtctgcaagctgcttatgctaatttgcatactgaccaagaacgtgattacttcatgcagcgttaccatgatgttatttcttcatttggaggtaaaacctcttatgacgctgacaaccgtcctttacttgtcatgcgctctaatctctgggcatctggctatgatgttgatggaactgaccaaacgtcgttaggccagttttctggtcgtgttcaacagacctataaacattctgtgccgcgtttctttgttcctgagcatggcactatgtttactcttgcgcttgttcgttttccgcctactgcgactaaagagattcagtaccttaacgctaaaggtgctttgacttataccgatattgctggcgaccctgttttgtatggcaacttgccgccgcgtgaaatttctatgaaggatgttttccgttctggtgattcgtctaagaagtttaagattgctgagggtcagtggtatcgttatgcgccttcgtatgtttctcctgcttatcaccttcttgaaggcttcccattcattcaggaaccgccttctggtgatttgcaagaacgcgtacttattcgccaccatgattatgaccagtgtttccagtccgttcagttgttgcagtggaatag |
| F3 WT | tattcgccaccatgattatgaccagtgtttccagtccgttcagttgttgcagtggaatagtcaggttaaatttaatgtgaccgtttatcgcaatctgccgaccactcgcgattcaatcatgacttcgtgataaaagattgagtgtgaggttataacgccgaagcggtaaaaattttaatttttgccgctgaggggttgaccaagcgaagcgcggtaggttttctgcttaggagtttaatc**atg**tttcagacttttatttctcgccataattcaaactttttttctgataagctggttctcacttctgttactccagcttcttcggcacctgttttacagacacctaaagctacatcgtcaacgttatattttgatagtttgacggttaatgctggtaatggtggttttcttcattgcattcagatggatacatctgtcaacgccgctaatcaggttgtttctgttggtgctgatattgcttttgatgccgaccctaaattttttgcctgtttggttcgctttgagtcttcttcggttccgactaccctcccgactgcctatgatgtttatcctttgaatggtcgccatgatggtggttattataccgtcaaggactgtgtgactattgacgtccttccccgtacgccgggcaataacgtttatgttggtttcatggtttggtctaactttaccgctactaaatgccgcggattggtttcgctgaatcaggttattaaagagattatttgtctccagccacttaagtgaggtgattt**atg**tttggtgctattgctggcggtattgcttctgctcttgctggtggcgccatgtctaaattgtttggaggcggtcaaaaagccgcctccggtggcattcaaggt |
| F3a (TAG) | tattcgccaccatgattatgaccagtgtttccagtccgttcagttgttgcagtggaatagtcaggttaaatttaatgtgaccgtttatcgcaatctgccgaccactcgcgattcaatcatgacttcgtgataaaagattgagtgtgaggttataacgccgaagcggtaaaaattttaatttttgccgctgaggggttgaccaagcgaagcgcggtaggttttctgcttaggagtttaatc**tag**tttcagacttttatttctcgccataattcaaactttttttctgataagctggttctcacttctgttactccagcttcttcggcacctgttttacagacacctaaagctacatcgtcaacgttatattttgatagtttgacggttaatgctggtaatggtggttttcttcattgcattcagatggatacatctgtcaacgccgctaatcaggttgtttctgttggtgctgatattgcttttgatgccgaccctaaattttttgcctgtttggttcgctttgagtcttcttcggttccgactaccctcccgactgcctatgatgtttatcctttgaatggtcgccatgatggtggttattataccgtcaaggactgtgtgactattgacgtccttccccgtacgccgggcaataacgtttatgttggtttcatggtttggtctaactttaccgctactaaatgccgcggattggtttcgctgaatcaggttattaaagagattatttgtctccagccacttaagtgaggtgattt**atg**tttggtgctattgctggcggtattgcttctgctcttgctggtggcgccatgtctaaattgtttggaggcggtcaaaaagccgcctccggtggcattcaaggt |
| F3b (TAG) | tattcgccaccatgattatgaccagtgtttccagtccgttcagttgttgcagtggaatagtcaggttaaatttaatgtgaccgtttatcgcaatctgccgaccactcgcgattcaatcatgacttcgtgataaaagattgagtgtgaggttataacgccgaagcggtaaaaattttaatttttgccgctgaggggttgaccaagcgaagcgcggtaggttttctgcttaggagtttaatc**atg**tttcagacttttatttctcgccataattcaaactttttttctgataagctggttctcacttctgttactccagcttcttcggcacctgttttacagacacctaaagctacatcgtcaacgttatattttgatagtttgacggttaatgctggtaatggtggttttcttcattgcattcagatggatacatctgtcaacgccgctaatcaggttgtttctgttggtgctgatattgcttttgatgccgaccctaaattttttgcctgtttggttcgctttgagtcttcttcggttccgactaccctcccgactgcctatgatgtttatcctttgaatggtcgccatgatggtggttattataccgtcaaggactgtgtgactattgacgtccttccccgtacgccgggcaataacgtttatgttggtttcatggtttggtctaactttaccgctactaaatgccgcggattggtttcgctgaatcaggttattaaagagattatttgtctccagccacttaagtgaggtgattt**tag**tttggtgctattgctggcggtattgcttctgctcttgctggtggcgccatgtctaaattgtttggaggcggtcaaaaagccgcctccggtggcattcaaggt |
| F4 | ggcgccatgtctaaattgtttggaggcggtcaaaaagccgcctccggtggcattcaaggtgatgtgcttgctaccgataacaatactgtaggcatgggtgatgctggtattaaatctgccattcaaggctctaatgttcctaaccctgatgaggccgcccctagttttgtttctggtgctatggctaaagctggtaaaggacttcttgaaggtacgttgcaggctggcacttctgccgtttctgataagttgcttgatttggttggacttggtggcaagtctgccgctgataaaggaaaggatactcgtgattatcttgctgctgcatttcctgagcttaatgcttgggagcgtgctggtgctgatgcttcctctgctggtatggttgacgccggatttgagaatcaaaaagagcttactaaaatgcaactggacaatcagaaagagattgccgagatgcaaaatgagactcaaaaagagattgctggcattcagtcggcgacttcacgccagaatacgaaagaccaggtatatgcacaaaatgagatgcttgcttatcaacagaaggagtctactgctcgcgttgcgtctattatggaaaacaccaatctttccaagcaacagcaggtttccgagattatgcgccaaatgcttactcaagctcaaacggctggtcagtattttaccaatgaccaaatcaaagaaatgactcgcaaggttagtgctgaggttgacttagttcatcagcaaacgcagaatcagcggtatggctcttctcatattggcgctactgcaaaggatatttctaatgtcgtcactgatgctgcttctggtgtggttgatatttttcatggtattgataaagctgttgccgatacttggaacaatttctggaaagacggtaaagctgatggtattggctctaatttgtctaggaaataaccgtcaggattgacaccctcccaattgtatgttttcatgcctccaaatcttggaggcttttttatggttcgttcttattacccttctgaatgtcacgctgattattttgactttgagcgtatcgaggctcttaaacctgctattgaggcttgtggcatttctactctttctcaatccccaatgcttggcttccataagcagatggataaccgcatcaagctcttggaagagattctgtcttttcgtatgcagggcgttgagttcgataatggtgatatgtatgttgacggccataaggctgcttctgacgttcgtgatgagtttgtatctgttactgagaagttaatggatgaattggcacaatgctacaatgtgctcccccaacttgatattaataacactatagaccaccgccccgaaggggacgaaaaatggtttttagagaacgagaagacggttacgcagttttgccgcaagctggctgctgaacgccctcttaaggatattcgcgatgagtataattaccccaaaaagaaaggtattaaggatgagtgttcaagattgctggaggcctccactatgaaatcgcgtagaggctttgctattcagcgtttgatgaatgcaatgcgacaggctcatgctgatggttggtttatcgtttttgacactctcacgttggctgacgaccgattagaggcgttttatgataatcccaatgctttgcgtgactattttcgtgatattggtcgtatggttcttgctgccgagggtcgcaaggctaatgattcacacgccgactgctatcagtatttttgtgtgcctgagtatggtacagctaatggccgtcttcatttccatgcggtgcactttatgcggacacttcctacaggtagcgttgaccctaattttggtcgtcgggtacgcaatcgccgccagttaaatagcttgcaaaatacgtggccttatggttacagtatgcccatcgcagttcgctacacgcaggacgctttttcacgttctggttggttgtggcctgttgatgctaaaggtgagccgcttaaagctaccagttatatggctgttggtttctatgtggctaaatacgttaacaaaaagtcagatatggaccttgctgctaaaggtctaggagctaaagaatggaacaactcactaaaaaccaagctgtcgctacttcccaagaagctgttcagaatcagaatgagccgcaacttcgggatgaaaatgctcacaatgacaaatctgtccacggagtgcttaatccaacttaccaagctgggttacgacgcgacgccgttcaaccagatattgaagcagaacgcaaaaagagagatgagattgaggctgggaaaagttactgtagccgacgttttggcggcgcaacctgtgacgacaaatctgctcaaatttatgcgcgcttcgataaaaatgattggcgtatccaacctgcagagttttatcgcttccatgacgcagaagttaacactttcggatatttctgatgagtcgaaaaattatcttgataaagcaggaattactactgcttgtttacgaattaaatcgaagtggactgctggcggaaaatgagaaaattcgacctatccttgcgcagctcgagaagctcttactttgcgacctttcgccatcaactaacgattctgtcaaaaactgacgcgttggatgaggagaagtggcttaatatgcttggcacgttcgtcaaggactggtttagatatgagtcacattttgttcatggtagagattctcttgttgacattttaaaagagcgtggattactatctgagtccgatgctgttcaaccactaataggtaagaaatcatgagtcaagttactgaacaatccgtacgtttccagaccgctttggcctctattaagctcattcaggcttctgccgttttggatttaaccgaagatgatttcgattttctgacgagtaacaaagtttggattgctactgaccgctctcgtgctcgtcgctgcgttgaggcttgcgtttatggtacgctggactttgtgggataccctcgctttcctgctcctgttgagtttattgctgccgtcattgcttattatgttcatcccgtcaacattcaaacggcctgtctcatcatggaaggcgctgaatttacggaaaacattattaatggcgtcgagcgtccggttaaagccgctgaattgttcgcgtttaccttgcgtgtacgcgcaggaaacactgacgttcttactgacgcagaagaaaacgtgcgtcaaaaattacgtgcggaaggagtgatgtaatgtctaaaggtaaaaaacgttctggcgctcgccctggtcgtccgcagccgttgcgaggtactaaaggcaagcgtaaaggcgctcgtctttggtatgtaggtggtcaacaattttaattgcaggggcttcggccccttacttgag |

**Table S2. Primers used to amplify WT φX174 genome for fragments F2 and F4.**

| **Identifier** | **Description** | **Sequence** |
| --- | --- | --- |
| 793 | Forward primer. PCR amplifies F4 backbone | GGCGCCATGTCTAAATTGTT |
| 794 | Reverse primer. PCR amplifies F4 backbone | CTCAAGTAAGGGCCGAAG |
| 795 | Forward primer. PCR amplifies F2 backbone | GAGGCTAACCCTAATGAGCTTA |
| 796 | Reverse primer. PCR amplifies F2 backbone | CTATTCCACTGCAACAACTGAA |

**Table S3. Primers used for Gibson Assembly of aiφX174 mutants.** F3a refers to F3 containing the amber initiator for the G gene. F3b refers to F3 containing the amber initiator for the H gene.

| **Identifier** | **Description** | **Sequence** |
| --- | --- | --- |
| 803 | F1 Fragment F | GAAGACCAGGAGCACCTGCtAGGTGGTCAACAATTTTAAT |
| 804 | F1 Fragment R | GAAGACCAAGCGCACCTGCCGAGCATCATCTTGATTAAG |
| 805 | F2 Fragment F | GAAGACCAGGAGCACCTGCAGATGATGCTCGTTATGGTT |
| 806 | F2 Fragment R | GAAGACCAAGCGCACCTGCGGAAACACTGGTCATAATCA |
| 807 | F3a Fragment F | GAAGACCAGGAGCACCTGCACCAGTGTTTCCAGTCCGTT |
| 808 | F3a Fragment R | GAAGACCAAGCGCACCTGCGCGTTGACAGATGTATCCATCT |
| 809 | F3b Fragment F | GAAGACCAGGAGCACCTGCATCTGTCAACGCCGCTAATC |
| 810 | F3b Fragment R | GAAGACCAAGCGCACCTGCACCGCCTCCAAACAATTTAGA |
| 811 | F4 Fragment F | GAAGACCAGGAGCACCTGCTTTGGAGGCGGTCAAAAAGC |
| 812 | F4 Fragment R | GAAGACCAAGCGCACCTGCGTTGACCACCTACATACCAAAG |
